# Sedimentary ancient DNA expands bioarchaeological evidence and interpretation at Roman Harelbeke

**DOI:** 10.64898/2026.11.06.764647

**Authors:** Kadir Toykan Özdoğan, Fabricio Furni, Kevin Nota, Jolijn Erven, Jordy Aal, Simone Reurings, Lars den Boef, Guillermo Sanchez, Roy van Mousch, G. Arjen de Groot, Gertjan Plets

**Affiliations:** Department of History and Art History, Utrecht University, Drift 6, Utrecht, 3512 BS, Netherlands; Animal Ecology, Wageningen Environmental Research, P.O. Box 47, Wageningen, Gelderland, 6700 AA, the Netherlands; Faculty of Science, Department of Earth Sciences, Geology and Geochemistry Cluster, Vrije Universiteit Amsterdam, Amsterdam, The Netherlands; Max Planck Institute for Evolutionary Anthropology, Leipzig, Germany; Groningen Institute of Archaeology, University of Groningen, Groningen 9712 ER, Netherlands; BAAC, 5222 BS ’s-Hertogenbosch, the Netherlands

## Abstract

Sedimentary ancient DNA (sedaDNA) can expand archaeological research by recovering broad biological signals from small amounts of sediment. Here, we apply shotgun sedaDNA metagenomics and target capture (also known as hybridization capture) to disused Roman wells from the vicus of Harelbeke, Belgium, dated to ca. 160-300 CE, and integrate the results with existing bioarchaeological datasets. Moving beyond individual introduced taxa as proxies for Roman influence, we reconstruct community-level profiles spanning consumption-related, synanthropic, and environmental organisms. SedaDNA reproduced most established bioarchaeological signals, including cultivated plants and domestic animals, while also recovering additional environmental taxa that established methods underrepresented. Mitochondrial target capture strengthened selected faunal detections and enabled phylogenetic placement of a pig (*Sus scrofa*) within the European mitochondrial lineage. The predominance of previously established consumption-related taxa is consistent with subsistence continuity and selective adoption of Roman-introduced resources, although taxon counts alone cannot establish their relative importance. Analytical-effort modeling further indicates that batch-based sedaDNA workflows can provide broad biological profiles at lower cost and time investment than established archaeological methods. Overall, sedaDNA complements conventional bioarchaeology by broadening taxonomic recovery, supporting scalable sampling, and providing a broad-spectrum approach to reconstructing settlement ecology, subsistence, and human-environment interaction.

## Introduction

In Roman-period archaeology, introduced plants and animals have long served as valuable indicators of connectivity, exchange, shifting consumption practices, and Roman cultural influence. Their significance is best understood when considered alongside locally established taxa and the wider biological communities present in settlement deposits (Lodwick & Rowan, 2022; van der Veen et al., 2008; Versluys, 2014). Rather than representing simple cultural replacement, introduced taxa can be interpreted as part of changing systems of movement, selective adoption, and local continuity, in which new species were incorporated alongside established foodways and regional ecological practices (Bakels & Jacomet, 2003; Bouby et al., 2022; van der Veen et al., 2008).

This broader perspective requires approaches that integrate findings related to consumption, animal management, synanthropic species (species associated with human settlements or activity), and environmental background within a single interpretive framework. Conventional archaeobotanical and archaeozoological analyses provide the foundation for such reconstructions by identifying taxa preserved as macroremains, microfossils, pollen, and bone, including both human-associated and environmental taxa. However, preservation, recovery practices, and analytical priorities shape these assemblages (Lodwick & Rowan, 2022), and identification and interpretation often focus most strongly on taxa with clearer archaeological relevance, such as cultivated plants and domestic animals. As a result, conventional datasets may underrepresent smaller, more fragile, or environmentally informative taxa.

Over the last two decades, the field of ancient DNA (aDNA) has grown and pushed the boundaries of aDNA recovery (Kjær et al., 2022; van der Valk et al., 2021, and for a full review of the field, see Orlando et al., 2021). SedaDNA has proven especially valuable for detecting biomolecular traces that may not be visible in preserved remains alone but are present in cave deposits, lake cores, and other sedimentary archives (Gelabert et al., 2021; Parducci et al., 2019; Vernot et al., 2021). From a single sample, sedaDNA can recover evidence of animal presence (Pedersen et al., 2016), synanthropic organisms (Søe et al., 2018), local vegetation (Parducci et al., 2017), and the broader ecological context (Wang et al., 2021), with less bias toward human-related taxa than some conventional bioarchaeological methods. This broader range of evidence is especially useful for examining taxa related to everyday socio-economic practices such as food consumption and animal use, which connect environment, economy, and culture. Furthermore, when DNA preservation allows or is combined with target capture, sedaDNA can also move beyond taxonomic detection by recovering DNA sequences suitable for population dynamics, providing a potential bridge between community-level reconstruction and questions of animal mobility, geographic origin or population turnovers (Gelabert et al., 2021; Pedersen et al., 2021; Vernot et al., 2021). Yet despite methodological advances, sedaDNA remains underutilized in archaeology, and especially in Roman archaeology (Aldeias & Stahlschmidt, 2024; Özdoğan et al., 2024).Our study provides a proof of concept using disused wells from the Roman Vicus of Harelbeke, Belgium, spanning two archaeological phases (Phase II, ca. 160 - 225 CE, and Phase III, ca. 225 - 300 CE). This setting is particularly suitable for testing the added value of sedaDNA since it combines a Roman-period settlement context, phased well deposits, and existing archaeobotanical and archaeozoological evidence, allowing molecular results to be interpreted against an established bioarchaeological baseline. Earlier archaeological excavations suggest these wells were likely reused as refuse/latrine contexts (Van Mousch, 2022). They therefore provide a concentrated archive for examining both settlement activity and environmental context (Deforce et al., 2021). These existing datasets document dominant domestic livestock and cultivated plants, while sedaDNA offers an opportunity to extend this picture to additional taxa.

Here, we employed sedaDNA metagenomics to reconstruct community-level taxonomic profiles spanning animals, invertebrates, and diverse plant groups, together with targeted enrichment of selected faunal taxa. We pursue four interconnected goals. First, we evaluate how sedaDNA complements conventional bioarchaeological methods by comparing taxonomic breadth and resolution while assessing the relative analytical effort and scalability of each approach. Second, we test whether target capture can strengthen faunal detections and, where preservation and comparative data permit, provide insight into mitochondrial lineage affiliation and possible geographic affinities. Third, we generate an integrated overview of faunal and floral communities at Roman Harelbeke by combining molecular and conventional evidence. Finally, we assess the relative representation of previously established and Roman-introduced consumption-related taxa to explore patterns consistent with continuity and selective adoption. Together, these aims provide a proof of principle for an integrated archaeological workflow in which sedaDNA complements specialist analyses while expanding Roman-period archaeology from individual food or indicator taxa toward broader reconstructions of settlement ecology, subsistence continuity, and human-environment interaction.

## Materials G Methods

### Site Information

Located in the Leie Valley (West Flanders, Belgium), the Roman settlement of Harelbeke was likely a *vicus*, a medium-sized village of approximately 200-300 inhabitants that served as a regional hub for trade, craft production, and civic life. Such settlements also functioned as colonial extraction nodes, facilitating the development of imperial infrastructure and exchange networks (Plets et al., 2013). Archaeological excavations since the 1960s have revealed a dense settlement with religious, economic, and potentially minor military significance, the latter suggested by second-century CE fortified structures. Established in the second half of the first century CE, the settlement peaked in the mid-second century before declining in the third century and eventually being abandoned. Its development was likely favored by its strategic position along Roman roads and proximity to forested marshlands, the wet river valley floor, and fertile farmland.

### Sampling and Lab Processes

During excavations at Kollegelaan, Harelbeke, in 2019, archaeologists discovered many well-preserved archaeological remains dating from the 2nd to the 3rd centuries. The team collected sediment cores from important wells and potential latrines for palynological research. Using these organic-rich sediment cores, sedaDNA samples were collected with sterile tools and contamination-minimizing equipment, taking care to avoid previously exposed surfaces during subsampling (Epp et al., 2019; Özdoğan et al., 2026). SedaDNA analysis was conducted on 31 sediment samples from six water wells and one pit feature (water wells: 501, 504, 510, 513, 515, 516; pit feature: S2213), spanning the period 160 CE to 300 CE. Supplementary materials provide detailed sample information (Supplementary Table 1).

DNA was extracted using the eDNA MagAttract PowerSoil Pro Kit (Qiagen GmbH, Germany), and sequencing libraries were prepared according to the protocols, including negative controls to assess lab contamination, as in Kircher et al. (2012) at the Centre for GeoGenetics, University of Copenhagen. Libraries were sequenced using 150-bp paired-end sequencing on an Illumina NovaSeq 6000, targeting approximately 50 million reads per sample.

The 18 samples with initial animal detections in the shotgun metagenomic data were selected for target capture. Only these samples were processed using two myBaits panels (Daicel Arbor Biosciences): a custom panel targeting 16 animal species mitochondrial genomes (Supplementary Table 2) and the predesigned plant panel Angiosperms-353.

Target capture was carried out following the manufacturer’s protocol (myBaits v5.0 Manual, Arbor Biosciences, 2020), with four to five libraries pooled per reaction (2 µL of each library (Seeber et al., 2024)). After pooling, the libraries underwent a 10-cycle pre-capture PCR to amplify target molecules (Armbrecht et al., 2021), followed by hybridization at 55°C for 24 hours (Murchie et al., 2021). Post-hybridization washes were performed at 65°C. We then performed a 21-cycle post-capture PCR to amplify the captured fragments (Rohland et al., 2022).

### Bioinformatics

#### Shotgun Metagenomics

A Holi-style exhaustive mapping approach was employed in the bioinformatic workflow, as described in Kjær et al. (2022), in short: beginning with adapter removal and read merging performed using AdapterRemoval v. 2.3.1 (Schubert et al., 2016), followed by dust filtering (preprocess command with a minimum read length of 30 bp) and duplicate removal using SGA v. 0.10 (Simpson & Durbin, 2012). The filtered reads were mapped against the whole nucleotide database (nt; release 268, split and indexed in 200 parts) from the National Center for Biotechnology Information (NCBI), and the PhyloNorway dataset split and indexed in seven parts (Alsos et al., 2020), using Bowtie 2 v. 2.4 -k 1000 (Langmead & Salzberg, 2012). Alignment files were merged (BAM format) with SAMtools v. 1.14 (Danecek et al., 2021). Taxonomic classification and damage profiling were conducted utilizing the metaDMG v. 0.38 workflow, which employed an LCA-based classification method (ngsLCA) with the following parameters: minimum significance of 2.0, a minimum similarity score of 0.95, a maximum similarity score of 1.0, a minimum mapping quality of 20, and Bayesian classification enabled (Michelsen et al., 2022). Afterward, post-classification filtering was implemented, retaining only taxa with a minimum damage percentage of 8% and at least 100 reads (Michelsen et al., 2022).

To account for uncertainty in species-level taxonomic assignments due to incomplete references, we reassigned all species-level detections obtained through shotgun metagenomics to their corresponding genera. Genus-level identifications were therefore used as the primary reporting resolution throughout the sedaDNA dataset. Where independent archaeological, archaeobotanical, archaeozoological, or ecological evidence supported species presence, we discussed them as plausible representatives of the detected genera. These interpretations considered known at the site, their regional distribution during the Roman period, and their ecological and historical plausibility. This distinction ensured that contextual knowledge could inform archaeological interpretation without overstating the taxonomic resolution supported by the molecular data.

### Target Capture

Target capture using the Angiosperms-353 panel yielded no usable plant sequence data and was therefore excluded from subsequent analyses. Mitochondrial target capture reads from 18 samples underwent adapter removal and merging using AdapterRemoval v2.3.1 (Schubert et al., 2016), followed by dust filtering with a minimum read length of 30 bp and duplicate removal using SGA v0.10 (Simpson & Durbin, 2012). Processed reads were analysed using Quicksand v2.6, retaining reads ≥35 bp (Szymanski et al., 2025). KrakenUniq v1.0.2 classified reads against the mitochondrial RefSeq release 221 database using 22-bp k-mers (Breitwieser et al., 2018). Families supported by ≥3 reads and ≥129 distinct k-mers were retained for alignment to pipeline-selected references using BWA with parameters -n 0.01 -o 2 -l 16500 (Li & Durbin, 2009). Alignments with mapping quality <25 were excluded, and alignment duplicates were removed. Coverage was calculated using SAMtools v1.15.1 (Danecek et al., 2021).

Quicksand’s default post-classification filters were applied, requiring a family sequence percentage ≥0.5% and a proportion of expected breadth ≥0.5, defined as observed breadth divided by expected breadth given the mean coverage (Szymanski et al., 2025). Reads assigned to the seven vertebrate families were then analyzed with kallisto v0.51.1 in single-end mode, using a 21-nt k-mer and family-matched mitochondrial reference panels (Bray et al., 2016). For each sample-family bin, the observed mean read length and standard deviation were supplied as fragment-length parameters. Kallisto estimated counts and TPM were summed across reference sequences assigned to the same genus using accession-verified taxonomy.

We assessed damage separately at the genus level, following the same approach Quicksand used for family-level damage assessment. We competitively mapped reads extracted from each family against downloaded reference mitogenomes, then removed duplicates and filtered low-complexity regions. Damage frequencies were then calculated using Quicksand’s damage assessment tool with double-stranded settings. Terminal 5′ C→T and 3′ G→A frequencies and their 95% confidence intervals were obtained from this analysis. We designated damage support as ++ or + when the lower confidence bound exceeded 9.5% at both or one terminus, respectively (Szymanski et al., 2025).

To assess the phylogenetic placement of the recovered pig aDNA, reads assigned to *Sus scrofa* from the sample UU0142 (well 513) were extracted from the Target Capture dataset and mapped to a circularized pig mitochondrial reference genome (Sscrofa 11.1; NC_000845.1), representing the European mitochondrial clade, using bwa aln with -l 1024, -n 0.01, and -o 2. BAM files were further deduplicated with Picard v2.25.2 using the MarkDuplicates command, with REMOVE_DUPLICATES = True and OPTICAL_DUPLICATE_PIXEL_DISTANCE = 2500. Reads were subsequently filtered to retain only mapped reads with a mapping quality (MQ) of at least 30, and 5 bp were trimmed from both ends to reduce the effects of terminal aDNA damage. SNPs were called using bcftools mpileup with -q 30 -Q 20 and -a FORMAT/AD,FORMAT/DP, followed by bcftools call -m with ploidy 2 (version 1.17). Although mitochondrial DNA is haploid, heterozygous positions were observed in the resulting VCF. These may reflect sequencing errors or residual post-mortem DNA damage, as commonly encountered in aDNA, but could also reflect the contribution of mitochondrial DNA from multiple individuals, as is possible in sedimentary aDNA (SedaDNA) (Vernot et al., 2021; Zavala et al., 2021). To investigate the nature of these polymorphisms, sites where the minor allele was supported by only one read were excluded. This filtering retained 41 polymorphic positions, eight of which had minor-allele coverage of at least three reads. These positions were distributed across the mitochondrial genome rather than concentrated in a particular region. To further minimize the potential influence of residual DNA damage, we also examined transversion polymorphisms only, retaining seven polymorphic positions. The presence of these polymorphisms, particularly transversions supported by multiple reads, suggests contributions from at least two pig individuals to the sediment sample, although this interpretation remains tentative.

To account for mixed alleles, we generated three alternative mitochondrial consensus sequences using different variant-calling and filtering approaches. First, diploid variant calls were filtered for a minimum depth of 2x. From this VCF, we generated two datasets: (approach 1) excluding all heterozygous sites, and (approach 2) retaining heterozygous sites with a minimum depth of 5x, and keep the major allele if it was supported by at least 70% of reads. Third (approach 3), variants were called using a haploid model and filtered for a minimum depth of 2x (changing -ploidy to 1 in previous command). For each of the three resulting VCFs, a major-allele consensus sequence was generated using bcftools consensus v1.17, with missing positions given as a bed file to be represented as N with the -m command.. Sequences were then decircularized by removing 15 bp from each end. The resulting consensus sequences were realigned with comparative modern and ancient pig mitochondrial genomes from previous studies (Bosse et al., 2014; Erven et al., 2025; Frantz et al., 2015, 2019; Ramírez et al., 2015) using mafft (Katoh & Standley, 2013). A phylogenetic tree was then reconstructed using RAxML (Stamatakis, 2014) raxmlHPC-PTHREADS -f a -m GTRGAMMA -p 45636 -x 45636 -# 1000 and visualized with figtreev1.5.0. All three consensus sequences consistently placed the sample within the European mitochondrial clade, regardless of the variant-calling and filtering approach used.

### Comparison of SedaDNA and Conventional Bioarchaeology Data

SedaDNA detections were compared with conventional archaeobotanical, palynological, and archaeozoological records from Harelbeke (Aal, 2022; Grabowski et al., 2022). Molecular detections were reported at genus level, while conventional identifications retained their reported species or genus resolution. Shotgun and target capture detections were combined into one molecular dataset. Since the datasets did not consistently represent the same deposits, comparisons focused on assemblage-level overlap. The unresolved sheep/goat record (*Ovis/Capra*) in the conventional bioarchaeology inventory was excluded from resolved taxon counts, neither genus was used for the comparison.

Taxa were assigned to three functional groups using a fixed genus-level lookup informed by the existing literature (Bakels, 2009; Groot, 2017; NDFF, 2026; Zeven, 1997): consumption-related taxa, comprising edible plants and animals associated with food procurement or husbandry; synanthropic taxa, comprising organisms associated with human settlements and disturbed habitats; and environmental taxa, reflecting surrounding vegetation and habitats. Genera whose constituent species spanned multiple functional groups were classified as functionally ambiguous: *Alopecurus, Avena, Capsella, Centaurea, Euphorbia, Linum, Papaver, Polygonum, Populus, Prunus, Sambucus*, and *Vicia*.

These genera were retained in the overall detection inventories but excluded from functional-group plots, percentage calculations, and consumption-related historical-status comparisons across all methods, including conventional species assigned to these genera. Ambiguity refers to functional classification, not uncertainty in taxonomic detection. *Homo* and taxa without a functional assignment were also excluded from the three-group comparison. The combined dataset retained conventional species identifications and additional molecular genera, removing redundant genus records where species from that genus were present. Functional proportions therefore describe detections at their reported resolution, not uniform genus richness, abundance, or dietary contribution.

Historical-status assignments used the supplied introduction-history lookup (Bakels, 2009; Groot, 2017; Zeven, 1997). Genus-level detections were classified as introduced only when all known species within the genus were considered introduced. Otherwise, their historical status remained unresolved, including cases where the introduction history was unknown.

### Analytical Effort Estimation

Conventional bioarchaeological costs were modeled per analyzed layer. Labor estimates, based on the professional experience of the co-authors, were 7-10 hours for archaeobotanical macroremains analysis and 5-8 hours for archaeozoology. Labor was valued at EUR 50 (rounded) per hour for both conventional bioarchaeological and sedaDNA analyses, based on average labor costs in the Netherlands (Statistics Netherlands, 2026). Palynology was estimated at EUR 500 per layer, based on quotations from three companies for a 20-sample project, including labor. Consumables were EUR 10 per layer for each method.

The sedaDNA model covered shotgun metagenomics only, using batches of 40 samples. Reagents and sequencing were estimated at EUR 300 per sample. Labor was modeled as 50-60 hours for the first batch, including laboratory work, bioinformatic setup, and analysis, and 20-30 hours for each additional batch. Batch numbers were rounded upward, and labor costs (EUR 50/hour) were divided by sample number before adding per-sample costs.

## Results

### SedaDNA-Based Results

#### Shotgun Metagenomics Detections

Overall, 23 of the 31 sampled sediments yielded meaningful results, with the richest assemblages occurring in several samples, particularly from wells 501, 504, 513, and 515. The initial shotgun assemblage comprised 139 genera, including *Homo* and 23 genera without a functional-group assignment. The comparative workflow additionally excluded five genera through its taxon-specific filtering rules, leaving 134 genera. Exclusion of *Homo* and the 18 remaining unassigned genera yielded 115 shotgun genera with an existing functional assignment. Excluding the 11 functionally ambiguous genera left 104 shotgun genera for functional-group analyses. The inventory included 12 animal genera, including *Homo*. Environmental taxa formed the largest functional group in the shotgun subset.

Most environmental detections belonged to the plant assemblage, including wetland and riparian genera such as *Carex*, *Juncus*, and *Salix*, alongside woodland- and grassland-associated genera such as *Alnus*, *Fraxinus*, *Phragmites*, *Menyanthes*, and *Thelypteris*. These genera include species characteristic of wetland, riparian, and woodland habitats, providing ecologically plausible interpretations of the molecular detections (NDFF, 2026). Consumption-related plant detections included genera associated with cereals, cultivated crops, and edible plants, such as *Hordeum, Secale, Triticum, Panicum, Rubus, Allium*, and *Beta*. Detected genera including *Papaver*, *Vicia, Prunus*, and *Sambucus* remained in the inventory but were classified as functionally ambiguous and excluded from consumption-related comparisons. Several of these genera include species documented in Roman-period agricultural and consumption practices, such as barley (*Hordeum vulgare*), rye (*Secale cereale*), millet (Panicum miliaceum), and beet (Beta vulgaris) (Bakels et al., 1997; Bakels & Jacomet, 2003). Although these species represent plausible interpretations in the archaeological context of Harelbeke, the molecular detections were retained at the genus level.

The faunal component was smaller but taxonomically diverse, comprising 12 animal genera, including *Homo*. The remaining animal genera were distributed across consumption-related, environmental, and synanthropic categories. Consumption-related detections included *Sus, Ovis, Gallus, Equus*, and *Clupea*. These genera include species documented in Roman-period animal husbandry, transport, or food consumption, such as pig/wild boar (*Sus scrofa*), sheep (*Ovis aries*), chicken (*Gallus gallus*), horse (Equus caballus), and Atlantic herring (*Clupea harengus*) (Groot, 2017; Van Neer et al., 2005). Their occurrence within the archaeological context is consistent with the presence of these animals. Environmental animal detections included *Apodemus*, *Arvicola*, *Rana*, and *Bradysia*, representing small mammals, amphibians, and insects potentially associated with the surrounding environment. Synanthropic detections included *Canis* and *Rattus*, consistent with the potential presence of dogs/wolves and rats within the settlement environment.

#### Target Capture and Mitogenome Analyses

Target capture of mitochondrial genomes recovered 55 genus-level signals across the 18 analyzed samples. These detections represented nine reference-derived genera: human (*Homo),* horse *(Equus),* pig *(Sus),* cattle *(Bos),* dog *(Canis),* sheep *(Ovis),* chicken *(Gallus),* and goose *(Anser)*. Relative to the shotgun assemblage, target capture added two genera: *Bos and Anser*. Wells 513 and 504 had the highest numbers and diversity of retained signals, with 20 signals across eight genera and 14 signals across six genera, respectively. Well 515 yielded ten detections across five genera, well 510 yielded five detections across five genera, and well 516 yielded four signals across four genera. Two sheep detections were recovered from well 501, whereas no signals passed the filters in the analyzed pit sample. Of the 55 retained detections, 38 showed positive deamination support (see Supplementary Figure 4).

Furthermore, the target capture enabled reconstruction of a nearly complete *Sus scrofa* mitochondrial genome. The strongest mitochondrial signal was recovered from well 513, where sample UU0142 reached approximately 5x mean mitogenome coverage and consistently clustered within the European mitochondrial clade (Figure 1, Supplementary Figure 5). Observed mitochondrial polymorphisms suggest contributions from multiple pig individuals, although distinct haplotypes could not be confidently reconstructed. The major consensus sequence of approach 1 was subsequently included in the phylogenetic tree (Figure 2), alongside sequences representing European *Sus scrofa* mitochondrial diversity. Within the European mitochondrial clade, UU0142 appears to cluster outside the group with Neolithic individuals from the Netherlands (Sw1, Sch1), Britain (DW3), and France (AA288); an Iron Age individual from Syria (AL613); a Roman-period individual from Hungary (AA409); a medieval individual from the Netherlands (KD024); and Norse-period individuals from the Faroe Islands (AA439, AA443, AA451), alongside modern Cinta Senese (CS01F02) and Chato Murciano (CM01F18) pigs (Figure 1, and see Supplementary Tables 3 & 4 for the full list of individuals added to the analysis). However, these relationships remain tentative because short internal branches and low bootstrap values limit resolution of its closest affinities.

**Figure 1.**
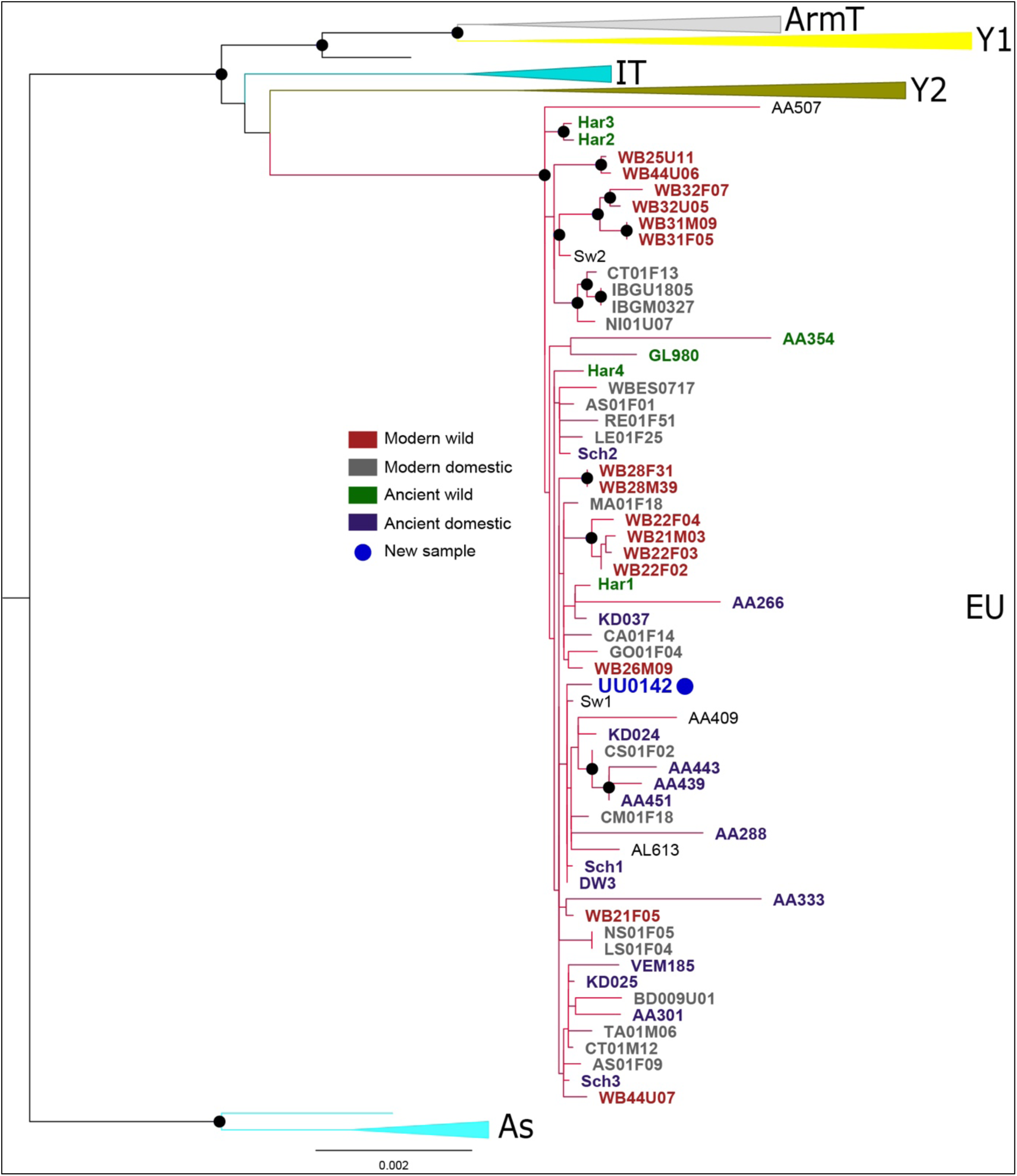
Maximum-likelihood tree of 115 mitogenomes from modern and ancient Sus scrofa for approach 1. This tree depicts the relationships between the common haplogroups found in Sus scrofa: Asia (AS, light blue), Anatolia/Southwest Asia (Y1, soft yellow), Armenia/Southwest Asia (Armt, gray), Southeast Europe (Y2, yellow green), Italy/South Europe (IT, cyan), and Europe (EU, dark red) (See Supplementary Figure 5 for non-collapsed version). Black circles indicate a bootstrap value >70.

**Figure 2.**
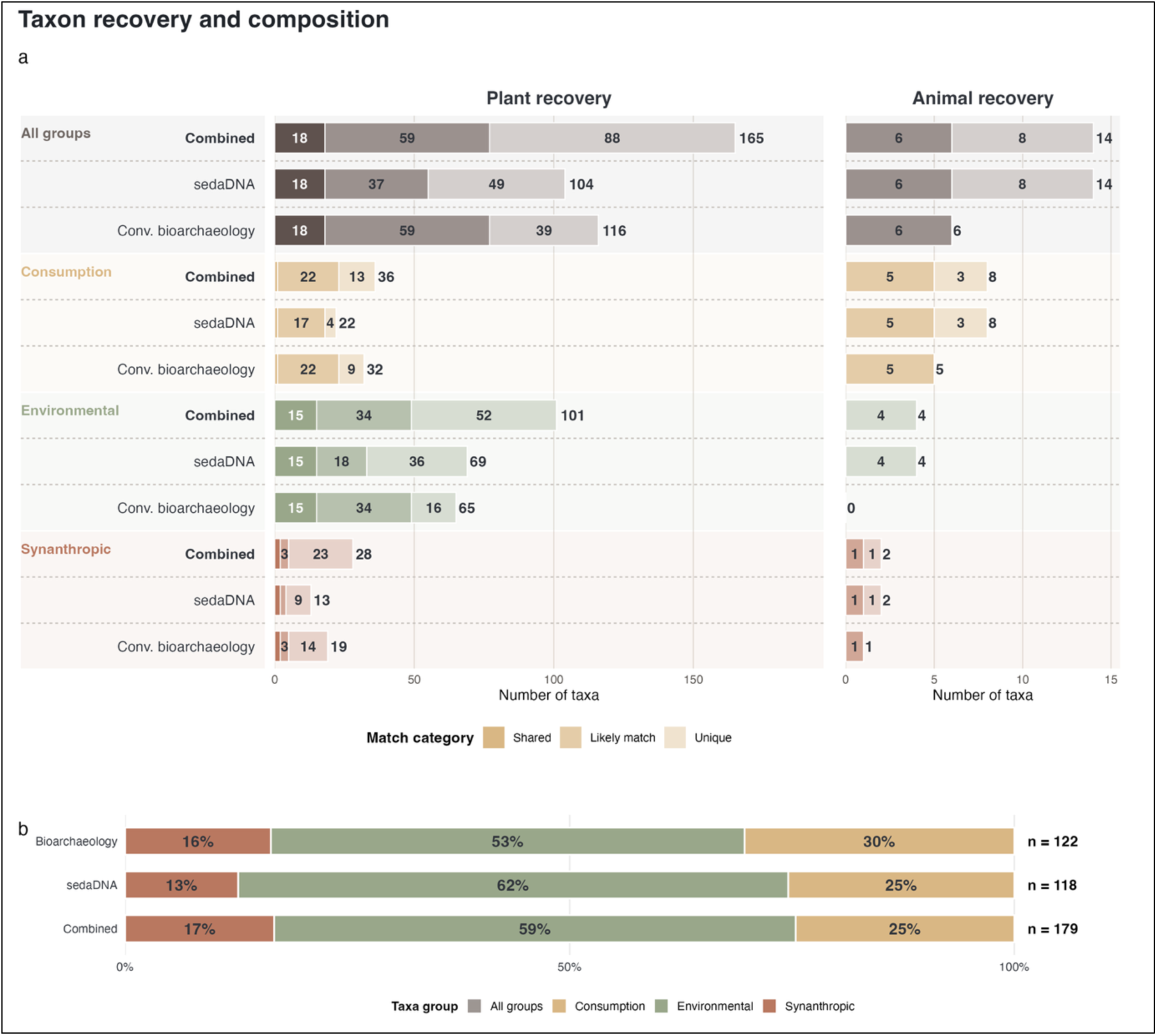
Recovery, method overlap, and functional group composition at reported taxonomic resolution. (a) Aligned plant and animal recovery bars show the combined dataset, sedaDNA, and conventional bioarchaeology within each functional group. Shades distinguish exact shared identifications, genus-level correspondences (likely matches), and method-specific detections (unique); numbers at bar ends indicate totals. (b) Functional group proportions within conventional bioarchaeology (122 taxa), sedaDNA (118 genera), and the combined dataset (179 taxa).

### Comparison with Conventional Bioarchaeological Evidence

The molecular analyses identified 118 genera, comprising 104 botanical and 14 animal taxa. Conventional bioarchaeological methods identified 122 taxa, comprising 116 botanical and 6 animal taxa, with the unresolved sheep/goat identification reported separately. Together, both approaches identified 179 taxa, of which 165 were assigned to plants and 14 to animals. (Figure 3).

**Figure 3.**
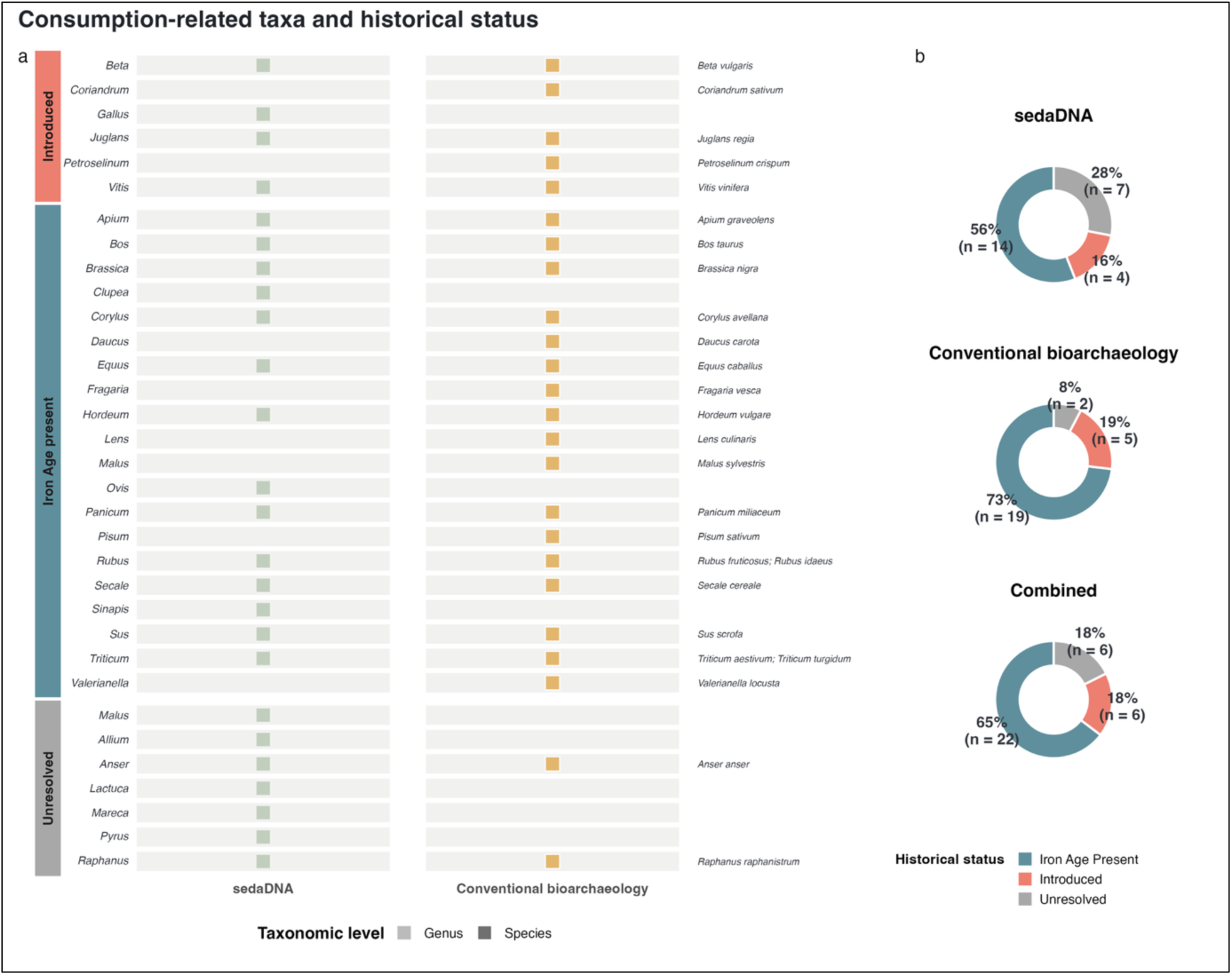
Consumption-related detections and historical status. (a) Genus-by-method display retaining conventional species identifications as annotations. Green and yellow squares identify molecular and conventional bioarchaeology detections; shading indicates the original identification resolution. (b) Donut charts summarize counted taxa at their reported resolution (sedaDNA, 25 genera; conventional bioarchaeology, 26 taxa; combined, 34 taxa). Blue, coral, and grey indicate Iron Age-present, introduced, and unresolved status.

Among plants, genus-level agreement encompassed cultivated and edible taxa, including cereals (*Hordeum*, *Secale*, *Triticum*, *Panicum*), fruit-bearing genera (*Prunus*, *Rubus*, *Sambucus*), and other consumption-related plants such as *Vitis*, *Corylus*, *Juglans*, and *Malus*. Conventional archaeobotanical analyses provided more specific identifications for several of these genera, including grape (*Vitis vinifera*), hazelnut (*Corylus avellana*), walnut (*Juglans regia*), and crab apple (*Malus sylvestris*). Correspondence also extended to environmental and synanthropic vegetation, including *Plantago*, *Betula*, *Salix*, *Quercus*, *Carex*, and *Juncus*. A comparable pattern was observed among animals, where shared genera such as *Sus*, *Bos*, *Anser*, *Equus*, and *Canis* were supported by both molecular and archaeozoological evidence. The sedaDNA detection of *Equus* also corresponded to the archaeozoological identification of horse (*Equus caballus*), illustrating how genus-level molecular signals can be placed within a more specific archaeological context.

Differences between the datasets further highlighted their complementary recovery characteristics. Conventional bioarchaeology provided species-level identifications for several cultivated plants and economically important animals that could not be resolved through shotgun sedaDNA. Archaeobotany uniquely identified taxa such as lentil (*Lens culinaris*), pea (*Pisum sativum*), coriander (*Coriandrum sativum*), and carrot (*Daucus carota*), while archaeozoology identified cattle (*Bos taurus*) and domestic goose (*Anser anser domesticus*), whose genera were not recovered through shotgun metagenomics. Mitochondrial target capture recovered *Bos* and *Anser*.

On the other hand, sedaDNA contributed a broader range of environmental genera, particularly among plants, including *Calluna*, *Frangula*, *Fraxinus*, *Phragmites*, and *Thelypteris*, as well as smaller vertebrates such as *Apodemus*, *Arvicola*, and *Rana*, which were not identified through conventional analyses. Additional animal detections included *Gallus* and *Clupea*, genera associated with consumption-related species such as chicken and herring. These differences reflect distinct preservation and method-specific recovery biases: conventional methods provide greater taxonomic specificity for many economically significant organisms, while sedaDNA captures a broader spectrum of environmental and less frequently represented taxa.

Environmental detections constituted 62% of the functionally classified sedaDNA subset (73/118), compared with 53% of the conventional bioarchaeological subset (65/122; Figure 3b). Consumption-related detections accounted for 25% (30/118) and 30% (37/122), respectively, while synanthropic detections accounted for 13% (15/118) and 16% (20/122). In the combined subset, environmental taxa comprised 59% (105/179), consumption-related taxa 25% (44/179), and synanthropic taxa 17% (30/179). These proportions exclude functionally ambiguous genera. Differences in reporting resolution contribute to these proportions, which represent taxonomic detections rather than organism abundance or dietary contribution.

### Archaeological Interpretation of the Combined Dataset

The The combined dataset contained 34 consumption-related taxa, representing 21.5% of the 158 detections included in functional comparisons (Figure 3b). Of these, 22 (64.7%) were assigned to Iron Age-present status, six (17.6%) to Roman-introduced status, and six (17.6%) remained unresolved (Figure 4b). Among the 28 taxa with a resolved status, 78.6% were Iron Age-present and 21.4% were Roman-introduced. These classifications describe historical occurrence rather than the geographic origin or dietary contribution of the recovered material. In the sedaDNA dataset, 14 of 25 consumption-related genera (56.0%) were classified as Iron Age-present, four (16.0%) as Roman-introduced, and seven (28.0%) remained unresolved. Conventional bioarchaeology recovered 19 Iron Age-present taxa (73.1%), five Roman-introduced taxa (19.2%), and two historically unresolved taxa (7.7%) among the 26 counted taxa.

**Figure 4.**
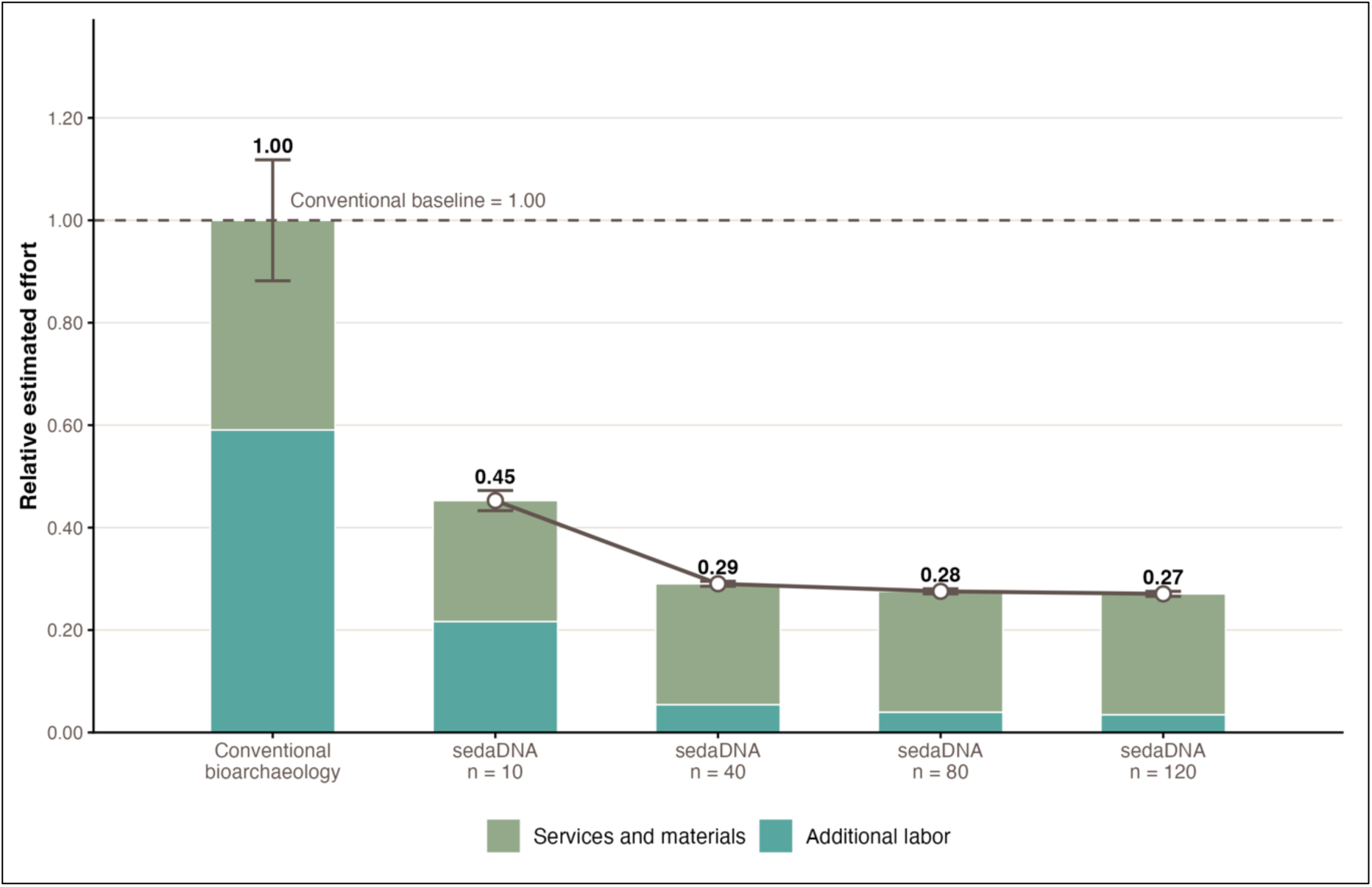
Relative modeled analytical effort for shotgun sedaDNA across increasing sample numbers, expressed as a cost-based proxy normalized to the conventional bioarchaeological midpoint (1.00). Stacked bars distinguish services and materials from additional labor, while error bars represent low- and high-labor scenarios rather than confidence intervals.

### Methodological Added Value of sedaDNA

Conventional bioarchaeological labor totaled 12-18 hours per layer. Consumables and palynology services totaled EUR 520 per layer. At EUR 50 per hour, conventional costs ranged from EUR 1,120-1,420 per layer, with a midpoint of EUR 1,270. The cost- based model placed shotgun sedaDNA below the combined conventional baseline at all sample sizes examined (Figure 4). The midpoint cost for shotgun metagenomics was EUR 575, EUR 368.75, EUR 350, and EUR 343.75 per sample for 10, 40, 80, and 120 samples, respectively, corresponding to relative values of 0.45, 0.29, 0.28, and 0.27 compared to conventional bioarchaeological analyses. Under these assumptions, declining per-sample shotgun costs were driven by distributing batch-level laboratory and bioinformatic labor across more samples.

## Discussion

### Complementarity Between sedaDNA and Conventional Bioarchaeological Methods

Our comparative results show substantial overlap between sedaDNA and conventional bioarchaeological datasets, supporting the reliability of the molecular identifications. By recovering signals also identified through established methods, including domesticated animals, cultivated crops, and synanthropic taxa, sedaDNA shows that it captures archaeologically meaningful patterns rather than only background environmental DNA. At the same time, the differences between datasets reflect the distinct taphonomic and analytical pathways through which each method records past biological communities.

Archaeobotanical and archaeozoological assemblages are shaped by preservation conditions, recovery strategies, and analytical focus (Lyman, 1994; Wright, 2005). Preservation conditions and recovery strategies influence which types and sizes of remains survive and are recovered, while research priorities influence which taxa researchers examine and report. Conventional methods provide direct morphological evidence for reconstructing crop use, livestock management, consumption practices, and other human activities. SedaDNA, by contrast, does not depend on the survival of identifiable macro-or micro-remains. It can therefore recover aDNA from organisms that are fragile, rare, widely dispersed in the environment, or unlikely to be preserved as visible archaeological remains (Brown et al., 2025; Özdoğan et al., 2024). These differences may contribute to the stronger representation of environmental taxa and the small mammals in the sedaDNA dataset, including taxa associated with woodland, wetland, riparian, and other local habitats, as well as rodents, such as *Arvicola* (water vole) and *Rattus* (rats).

The differences observed between methods should therefore not be interpreted simply as inconsistencies. Cases in which one method identifies a taxon at the genus level and another at the species level show that multi-proxy agreement does not require identical taxonomic precision. Instead, overlap at broader ecological, functional, or cultural levels may be equally meaningful. Method-specific detections also underline the risks of relying on a single proxy. The absence of small vertebrates and other environmentally sensitive taxa from the archaeozoological record likely reflects method-specific recovery and analytical bias, including the lack of systematic sieving, rather than their absence from the past environment (Shaffer & Sanchez, 1994). In contrast, the stronger representation of cultivated and consumption-related plants in archaeobotanical data reflects the value of macroremains for identifying taxa directly associated with food production, processing, and use.

Together, these results indicate that the methodological value of sedaDNA lies in the combination of biological breadth, low sediment requirement, and scalable batch processing. SedaDNA should therefore be viewed not as a replacement for conventional bioarchaeology, but as an efficient broad-spectrum baseline that can guide where more specialized archaeobotanical, archaeozoological, or targeted aDNA analyses are most useful.

#### Toward an Integrated Archaeological sedaDNA Workflow

The effort model indicates lower resource requirements for the modeled shotgun metagenomics workflow than for the combined conventional bioarchaeology workflow across the sample sizes examined. This finding depends on the specified costs, labor estimates, batch sizes, and analytical scope. SedaDNA can also serve as a screening tool for identifying deposits with high interpretive potential, especially where visible biological remains are rare, poorly preserved, or difficult to sample systematically.

Furthermore, the target capture results show how sedaDNA can move from broad biodiversity detection toward more focused genetic interpretation (Gelabert et al., 2025; Oberreiter et al., 2024). By increasing reference breadth and mean coverage, mitochondrial target capture strengthened some low-depth detections and enabled examination of selected taxa beyond presence or absence (Murchie et al., 2022; Vernot et al., 2021; Zavala et al., 2021).

The pig mitogenome recovered from well 513 demonstrates the potential of targeted capture of sedaDNA to identify mitochondrial lineages. The detected European mitochondrial lineage was already established in domestic pigs long before the Roman period (Larson et al., 2007), making a local or regional origin plausible. However, its broad geographic distribution also allows for introduction from elsewhere. Therefore, mitochondrial affiliation alone cannot establish the animal’s origin or demonstrate local continuity or individual mobility.

Several limitations therefore remain central to archaeological sedaDNA interpretation. SedaDNA does not detect all biological evidence equally well, particularly when DNA is highly degraded or present at very low endogenous levels. Read counts cannot be translated directly into organismal abundance or biomass without accounting for biological, depositional, preservation, and analytical biases (Chen & Ficetola, 2020). Species-level assignments may remain uncertain when reference databases are incomplete or contaminated (Oskolkov et al., 2025). Additionally, although researchers can distinguish aDNA from modern contamination, it remains a persistent concern even though they consistently apply contamination-aware sampling, laboratory procedures, authentication criteria, and negative controls (Özdoğan et al., 2026). These considerations are especially important for target capture-based population analyses, which require robust ancient DNA authentication, sufficient data across informative loci, and appropriate comparative datasets (Vernot et al., 2021).

An integrated archaeological sedaDNA workflow chould therefore be organized in tiers. Shotgun metagenomics can first be used to establish broad biological profiles across features and layers (Gelabert et al., 2021). Mitochondrial or chloroplast target capture can then strengthen detection of selected taxa and, where possible, recover lineage-level information (Murchie et al., 2022; Zavala et al., 2021). Finally, where preservation conditions and research questions justify the additional effort, nuclear target capture could be applied to investigate population structure, ancestry, or genetic affinities at higher resolution (Vernot et al., 2021).

### Dietary Practices and Cultural Continuity

The dominance of previously established (Iron Age) consumption-related taxa reflects continuity in established foodways, while Roman-introduced resources suggest selective adoption within wider exchange networks. This pattern aligns with conventional bioarchaeological evidence from Harelbeke (Aal, 2022; Grabowski et al., 2022), although mixed-resolution taxon counts do not directly demonstrate continuity in dietary proportions, local production, or subsistence practices.

Studies of Roman period archaeobotanical assemblages and frontier regions describe the incorporation of introduced foods alongside established resources (Bakels & Jacomet, 2003; Bouby et al., 2022; Groot, 2008; van der Veen et al., 2008). The Harelbeke assemblage offers a local case study for considering such selective adoption within wider exchange networks (Livarda & Orengo, 2015). This interpretation is consistent with approaches to Roman-period cultural change that emphasize selective integration. Traditional models of Romanization have been criticized for framing “Roman” and “local” as opposing categories and cultural change as a unidirectional process of assimilation (van Dommelen, 2014; Mattingly, 2004). Such perspectives risk obscuring regional diversity and the persistence of local practices, particularly in frontier contexts where cultural interaction was uneven and negotiated (Woolf, 2014). Approaches emphasizing hybridity, connectivity, and circulation instead conceptualize the Roman world as a landscape of “objects in motion,” in which interaction among people, goods, and practices generated new social configurations (Versluys, 2014).

Although directly comparable pre-Roman evidence from Harelbeke is needed to assess changes in resource use over time, combined sedaDNA and conventional bioarchaeological evidence provides a broader picture of established and introduced resources at Roman Harelbeke. These complementary results contribute to earlier discussions of how introduced resources were selectively integrated into existing local practices (Bouby et al., 2022; van der Veen et al., 2008; Versluys, 2014).

## Author Contributions

K.T. Özdoğan, G.A. de Groot, and G. Plets designed the study. K.T. Özdoğan conducted sampling, wrote the manuscript, performed bioinformatics analyses, and prepared Figures 1-4. G.A. de Groot and G. Plets contributed additions and suggestions throughout the manuscript and share senior authorship. F. Furni performed bioinformatics analyses and provided suggestions for the manuscript. K. Nota provided suggestions for data analyses, and J. Erven performed additional bioinformatics analyses. G. Sanchez helped design the Target Capture kit and provided input on the zooarchaeological focus. R. von Mousch provided samples and wrote the site description. S. Reurings, J. Aal, and L. den Boef performed bioarchaeological analyses and suggested revisions to the manuscript. All authors reviewed and approved the final version of the manuscript.

## Competing Interests

The authors declare no competing interests.

## Acknowledgments and Funding

We would like to thank the members of the C-LIMES project and the associated institutions for their valuable discussions and feedback during the development of this article. We also thank Xihan Chen, Kurt Kjær, and Ivo Laros for their assistance with sample processing and data production. This article derives from the project “Constructing the Limes: Employing citizen science to understand borders and border systems from the Roman period until today” (C-LIMES), funded by the Dutch Research Council (NWO) as part of the Dutch Research Agenda (2021–2026; project number NWA.1292.19.364).

## Declaration of Generative AI and AI-Assisted Technologies in the Writing Process

During the preparation of this manuscript, the authors used Grammarly for language editing and ChatGPT for assistance with clarity, readability, and plotting scripts. The authors reviewed and edited the resulting text, checked the data summaries and scripts, and took full responsibility for the manuscript’s content.

## Data and Code Availability

Raw sequencing reads are available on the European Nucleotide Archive (ENA) and can be found under the BioProject number: PRJEB129085.

## Supplementary Tables G Figures

**Supplementary Table 1**: Detailed sedaDNA sample information. See sheet S1_Sediment_samples in Supplementary_tables_and_data.xlsx.

**Supplementary Table 2**: Selected animal taxa and references for the Target Capture bait panel. See sheet S2_Capture_baits in Supplementary_tables_and_data.xlsx.

**Supplementary Table 3**: Genomic and geographical information of modern pig/wild boar comparison individuals. See sheet S3_Modern_pig_references in Supplementary_tables_and_data.xlsx.

**Supplementary Table 4**: Archaeological and genomic information of ancient pig/wild boar comparison individuals. See sheet S4_Ancient_pig_references in Supplementary_tables_and_data.xlsx.

**Supplementary Table 5**: The 55 mitochondrial reference sequences used in the separate genome-level mapping analysis, including accession versions, original taxon labels, sequence lengths, SHA-256 checksums, NCBI record URLs, and original FASTA headers. The panel totals 914,391 bp. See sheet S5_mtDNA_reference_panel in Supplementary_tables_and_data.xlsx.

**Supplementary Dataset 1**. Molecular genus inventories, conventional bioarchaeology species and genus identifications, counted mixed-resolution taxa, functional proportions, historical-status assignments, and functional-ambiguity flags and excluded records. Conventional sheep/goat remains unresolved and is listed separately. These data are provided in the D1_ sheets of Supplementary_tables_and_data.xlsx. The G_ sheets archive the earlier genus-only calculations before exclusion of functionally ambiguous genera; the Contents and Notes sheets distinguish these from the main comparisons. The workbook also includes Supplementary Tables 1-5, cost-model summaries, and display data for Figures 3-5.

---

**Supplementary Figure 1.**
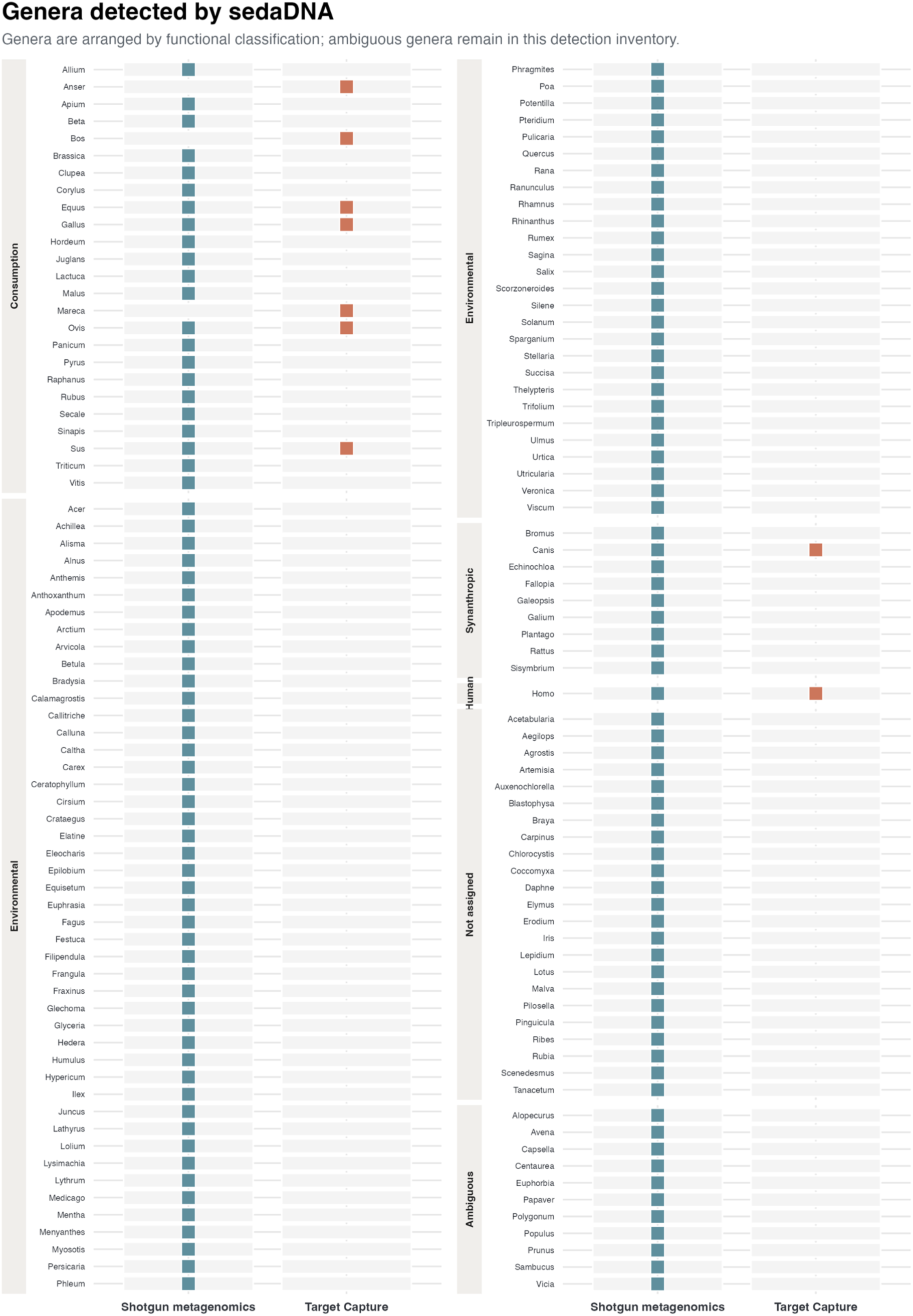
Genera detected by shotgun metagenomics and Target Capture, arranged alphabetically within functional classifications, including a separate ambiguous category. Squares indicate detection by each method. The union contains 142 genera: 139 in the initial shotgun inventory and nine in Target Capture, with Bos, Anser, and Mareca added by Target Capture. Homo, unassigned genera, and the 11 functionally ambiguous genera are shown in the inventory but excluded from the main functional comparisons. Five initial shotgun genera were additionally excluded by the comparative workflow; inclusion flags are provided in Supplementary Dataset 1.

**Supplementary Figure 2.**
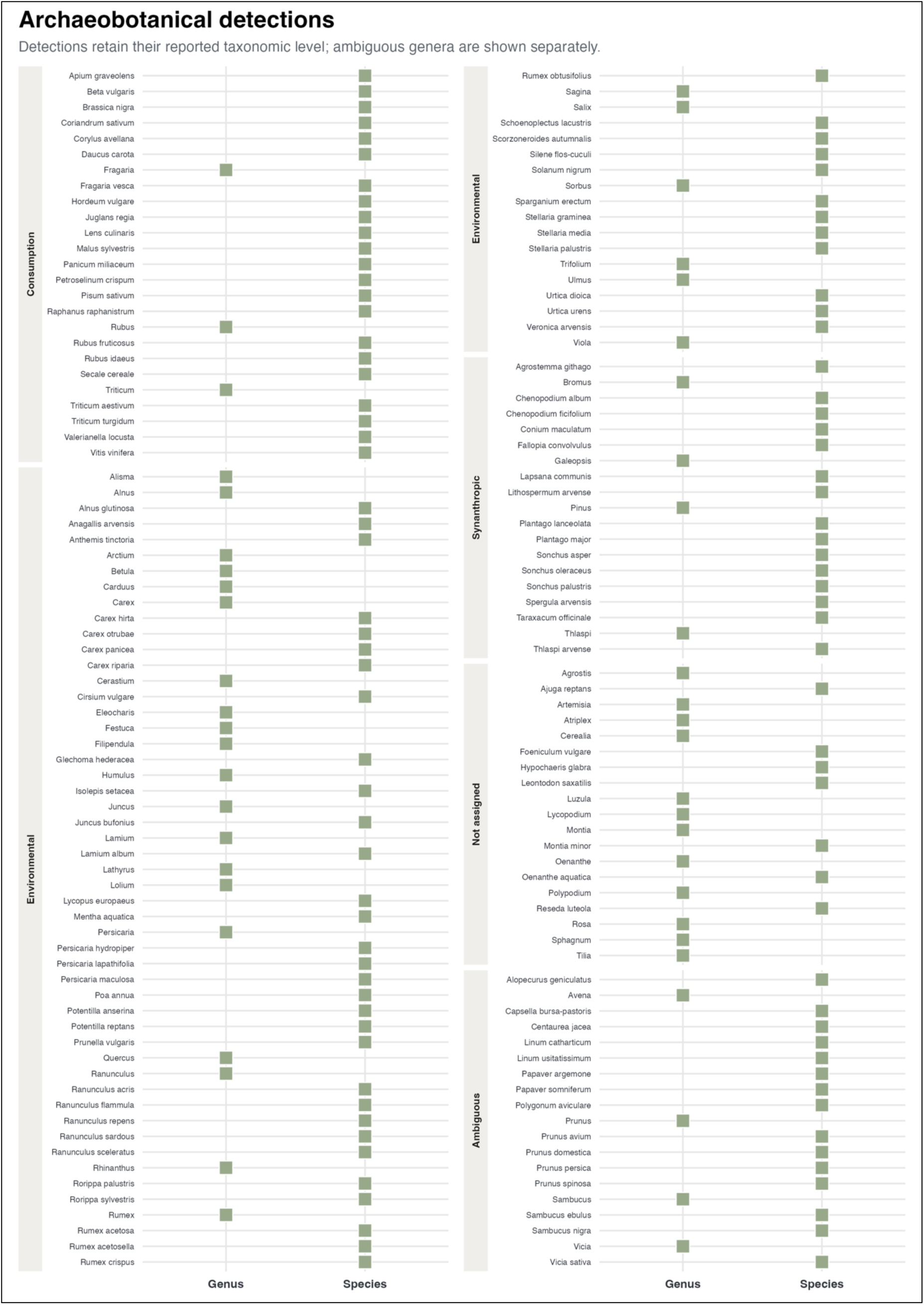
Archaeobotanical (including palynological) detection inventory, retaining functionally ambiguous genera in a separate category. Each square indicates whether the identification was reported at genus or species level. Ambiguous genera are excluded from the functional and consumption-related comparisons in Figures 3 and 4.

**Supplementary Figure 3.**
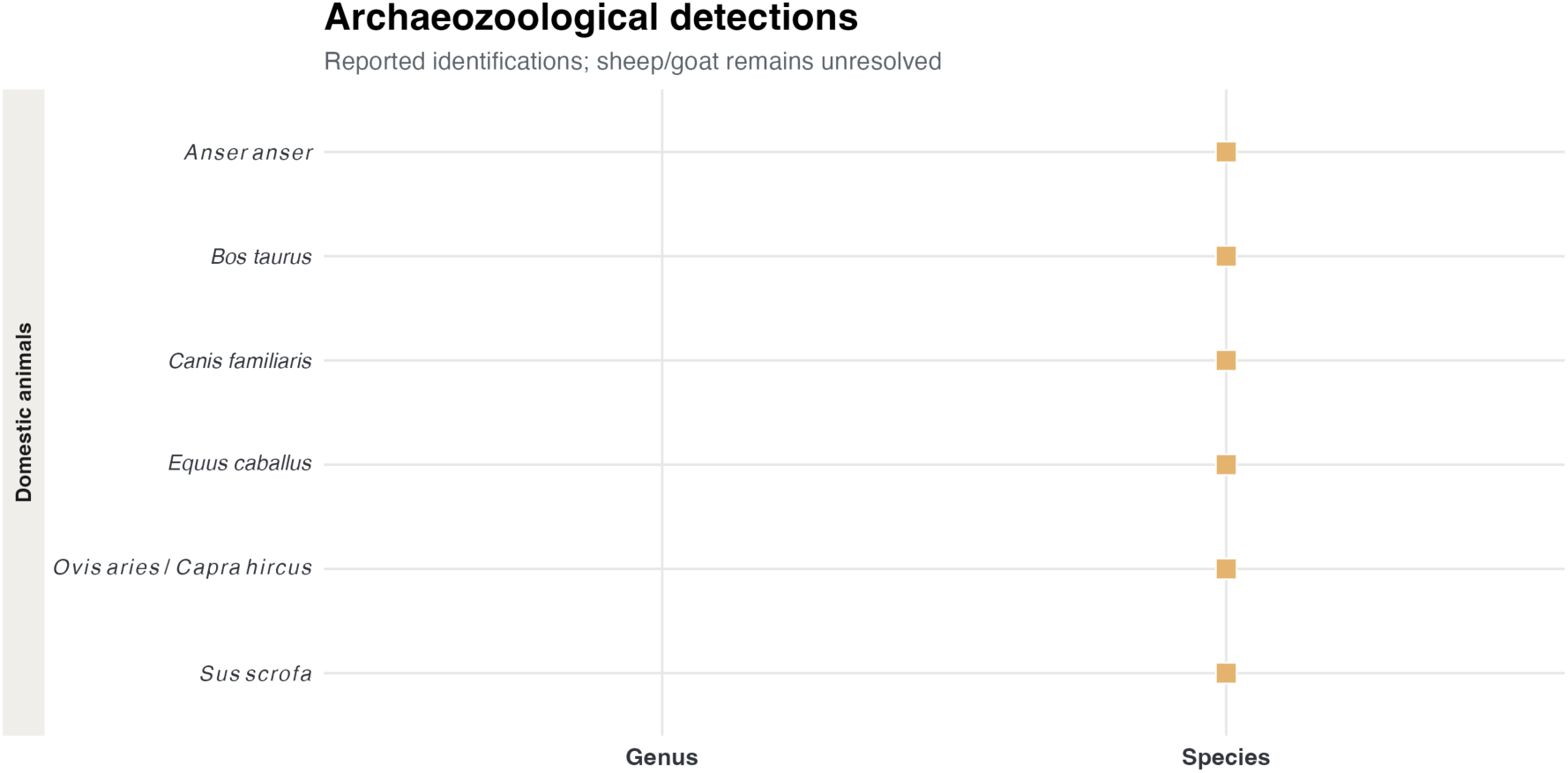
Conventional archaeozoological identifications. The sheep/goat record remains unresolved as *Ovis aries* / *Capra hircus* and is not counted as a confirmed *Ovis* or *Capra* detection in the resolved taxon comparisons. Domestic goose and dog labels follow the manuscript’s conventional bioarchaeology identification summary.

**Supplementary Figure 4.**
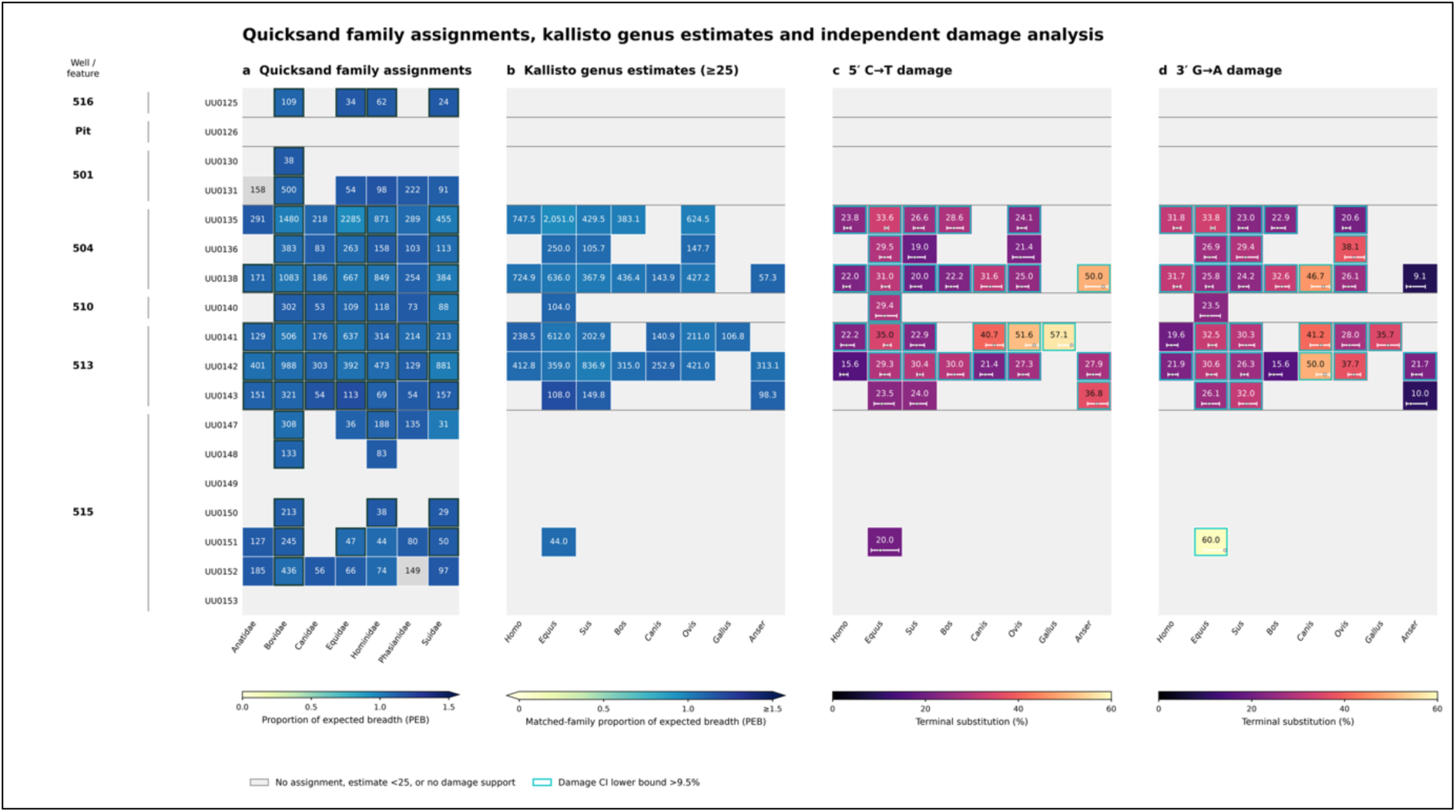
Family-level detection support, genus-level estimates, and ancient DNA damage patterns in mitochondrial target-capture data from 18 Harelbeke sediment samples. Rows represent samples grouped by archaeological well or pit feature. (a) Quicksand family assignments. Cell labels show the number of reads extracted, and fill color shows the proportion of expected breadth (PEB). Dark outlines mark sample-family bins passing the Quicksand post-classification thresholds (PSF ≥0.5% and PEB ≥0.5) that were used as Kallisto inputs; gray cells indicate an assignment for which PEB was unavailable, and blank cells indicate no reported family assignment. (b) Kallisto genus estimates, shown when estimated counts are ≥25 and the corresponding matched-family damage result is supported (+ or ++). Cell labels are Kallisto-estimated counts, not unique mapped-read counts; fill color shows PEB for the corresponding Quicksand family bin. (c,d) Terminal 5′ C→T and 3′ G→A substitution frequencies, respectively, calculated with Quicksand damage statistics on alignments from a separate competitive mapping of family-subset reads to species-level references. The damage values therefore describe the competitive-mapping genus groups. Miniature bars show 95% confidence intervals on a 0-60% scale; intervals extending beyond that range are clipped, while the substitution frequency is shown numerically. Cyan outlines mark cells where the lower confidence bound exceeds 9.5% at that terminus. Color gradients represent PEB in panels (a,b) and terminal substitution frequency in panels (c,d). Gray cells in (b-d) indicate no displayed result under the applied count and damage-support criteria; this does not establish biological absence.

**Supplementary Figure 5.**
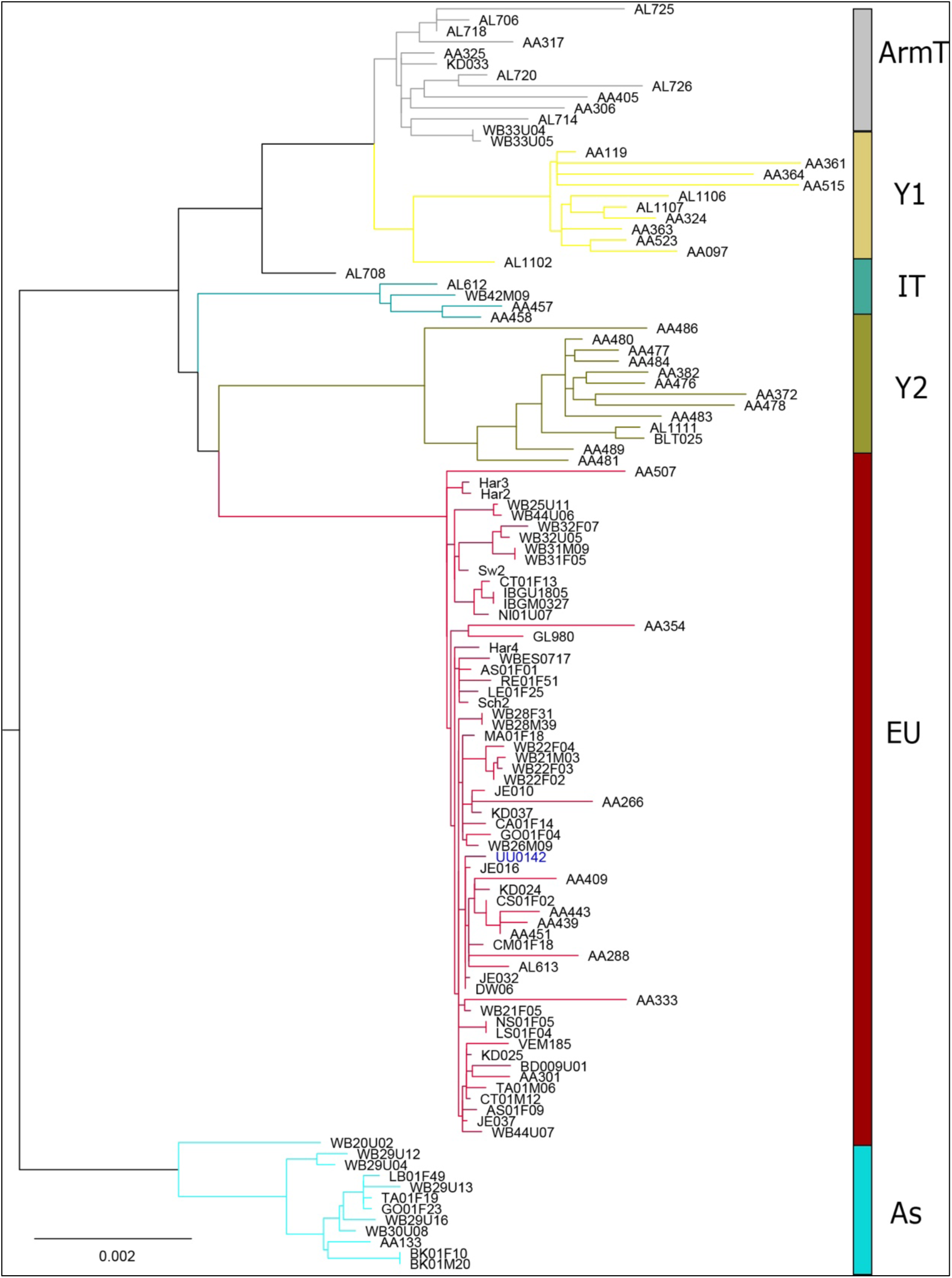
Maximum-likelihood phylogenetic tree of 115 modern and ancient *Sus scrofa* mitogenomes. The non-collapsed tree depicts relationships among the major mitochondrial haplogroups: Asia (AS, light blue), Anatolia/Southwest Asia (Y1, soft yellow), Armenia/Southwest Asia (ArmT, grey), Southeast Europe (Y2, yellow-green), Italy/Southern Europe (IT, cyan), and Europe (EU, dark red). The Harelbeke sample UU0142 is highlighted in blue and clusters within the European haplogroup. Branch lengths represent nucleotide substitutions per site; the scale bar indicates 0.002 substitutions per site.

## Supplementary Information

### Site and context description

Since the 1960s, numerous Roman-period finds have been uncovered in the Collegewijk district. These include the favisa containing numerous votive figurines on Olmenlaan, the impressive bronze cauldron discovered in the shaft of a well during the construction of the Sint-Amandscollege, and the large quantities of pottery and metal objects found throughout the area. In addition, several larger archaeological investigations have taken place between Dennenlaan and Kollegelaan, ranging from trial trenches and excavations to rescue excavations. These investigations have either not been fully elaborated, have been treated only in a very basic way, or have been highly selective due to the time and resources available. An overall synthesis is lacking but is currently in preparation. What the various investigations share is that they have not yet clearly identified the character of the settlement, which displayed considerable wealth. Both the excavations in 2010 and 2011 (Deconynck 2014) and in 2019 (Van Mousch 2022) revealed numerous traces of occupation belonging to a settlement that appears to have begun in the last quarter of the first century AD. From the second half of the second century onward, the occupation can probably be placed in its heyday, although the settlement may have continued to exist well into the third century with a certain degree of prosperity.

The 2019 excavation, from which the sedaDNA samples have been taken, showed that the settlement was located along (a crossroads of) roads, with its layout organized within several ditched enclosure blocks. Over time, these blocks shifted slightly. A notable modification took place in the last quarter of the second century (Phase II): along the roads, the enclosure blocks were provided with palisades, possibly with a raised walkway behind them and perhaps storage units or stable boxes. The settlement thus took on a much more defensive character. These rows of posts along the ditches and sandy roads were also identified during the 2010 and 2011 investigations.

Within the 2019 excavation area, the enclosure blocks were laid out as residential farmsteads, featuring houses, outbuildings, and water wells. Based on the wells, a broad occupational phasing can be established (Phases I-III). Particularly striking are the numerous and remarkable deposits found in the wells, which likely relate to the construction and abandonment of the wells. The buildings, from which little pottery was recovered, are difficult to phase. In terms of their location, they could fit into virtually all phases of occupation. In the southern part of the excavation area, feature density increases, yet only a few building structures could be reconstructed there. Nevertheless, several phases of occupation succeeded one another in this area.

A particularly notable phenomenon is present in one of the enclosure blocks. Within the ditched (and at a certain point fenced) area, rectangular and square pits were discovered in addition to building structures and wells, the precise function of which remains unknown. The walls of the pits appear to have been lined with wood or other organic material, which may indicate their use as storage facilities. The pits show a highly stratified fill, often with a strongly humic layer at the bottom. Selective archaeobotanical research has demonstrated that animal manure at least ended up in them. These features have not been used as latrines. Perhaps the pits primarily served as storage, with a secondary function as refuse or manure pits, but researchers have not yet been able to provide a definitive answer.

In contrast, after their abandonment, the wells appear to have been used as latrines. In the only well whose fill has undergone archaeobotanical analysis, indicators of human feces were identified. The six wells sampled for sedaDNA had not undergone archaeobotanical analysis.

### Conventional Archaeological Analyses

The Harelbeke faunal assemblage consists of 530 hand-collected animal remains, of which 374 were analyzed. The archaeozoological analyses indicate a dominance of domestic species, including *Bos taurus* (cattle), *Ovis aries*/*Capra hircus* (sheep/goat), *Sus scrofa domesticus* (pig), *Equus caballus* (horse), *Canis familiaris* (dog), and *Anser anser domesticus* (domestic goose). A relatively small number of mammalians, avian, and fish remains could not be identified to species level (Aal, 2022). Most identified species were also present in the various Roman wells (Supplementary Figure 3) and can therefore be considered representative of the larger assemblage.

The botanical records were based on 5 liters of soil samples taken from the wells and layered pits, which have a strongly humic bottom layer. From one well, both a botanical macrofossil and a pollen sample were studied. The macrofossil samples were wet sieved through a nest of sieves ranging from 2mm to 0.25mm. The fossils were handpicked from the residue using a stereomicroscope. The botanical remains show a diverse diet of several crops, some of which were already introduced before the Roman period, such as barley (*Hordeum vulgare*) and broomcorn millet (*Panicum miliaceum*), but others were Roman introductions that adapted to the local climate, like plum (*Prunus domestica*) and walnut (*Juglans regia*). Fruits such as grapes were probably imported, but could, on rare occasions, grow locally, while the pit of a peach shows that luxury goods were imported.

Alongside crops, the botanical macrofossils indicate local vegetation composed of plants well adapted to a highly disturbed, ruderal environment, such as that of a rural settlement. The pollen record indicates an open landscape with little to no woodland in the surrounding area, but occasional trees, such as hazel (*Corylus*), *are* present. (Grabowski et al., 2022)

### Target Capture Efficiency

Mitochondrial genome recovery efficiency was evaluated separately using a custom panel of 55 animal mitochondrial reference sequences, including a human reference, totaling 914,391 bp. Individual reference lengths ranged from 15,241 to 17,865 bp. The panel encompassed mammals, birds, fishes, amphibians and invertebrates, including domestic animals, wildlife taxa and groups of closely related species. Alignment against the combined panel allowed competitive mapping among the included references. Accession versions, reference lengths and original taxon labels are provided in Supplementary Table 5.

For genome-level mapping, we used filtered and deduplicated target capture FASTQ files before applying additional terminal trimming for marker screening. Reads were aligned against the combined mitochondrial panel using BWA-MEM v0.7.19-r1273 (bwa mem -t 8). Alignments with mapping quality below 30 were excluded using SAMtools v1.22 (samtools view -b -q 30), and retained alignments were coordinate-sorted and indexed. Per-position depth was calculated using samtools depth -a. For each sample and reference sequence, mean coverage was calculated as the summed depth divided by the number of reported reference positions, and breadth as the proportion of reported positions covered by at least one read.

**Supplementary Figure 6.**
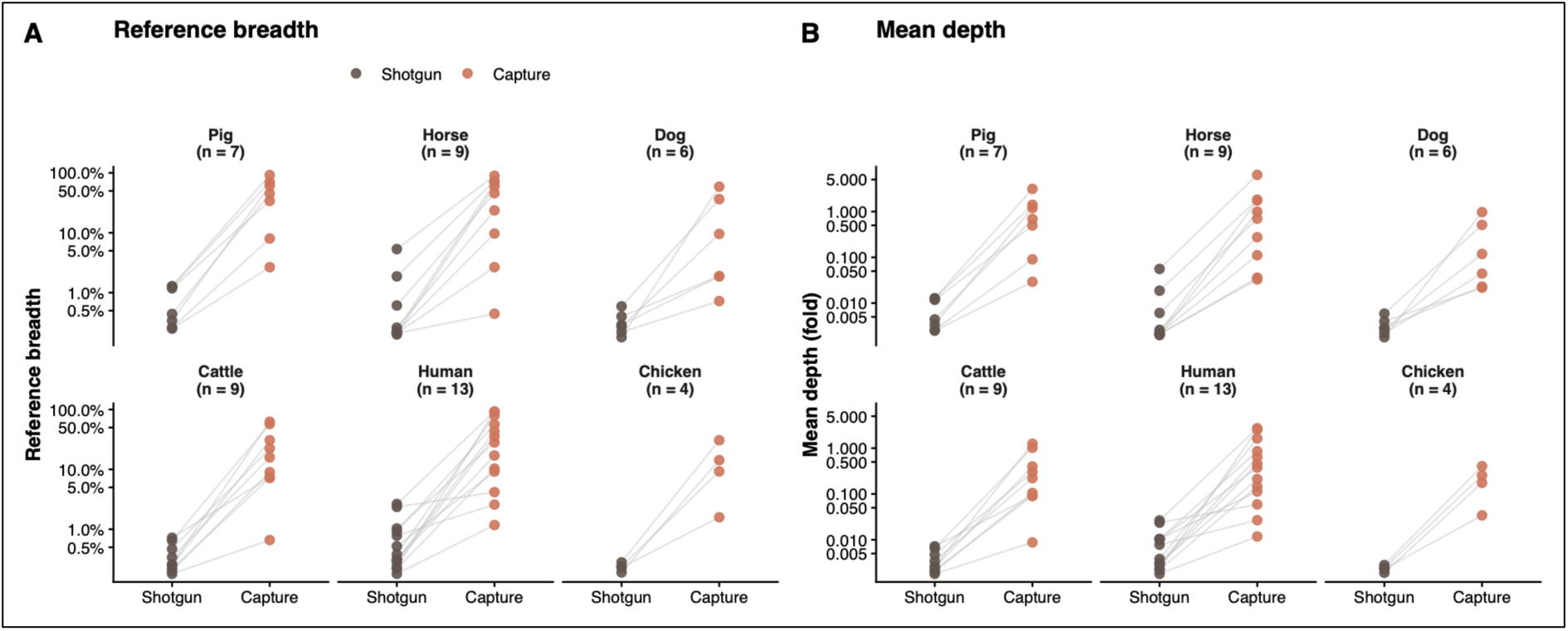
a) Reference breadth and b) mean depth for paired sample-reference detections recovered by both methods. Lines connect identical sample-reference combinations, with values shown on a log scale.

The comparison between shotgun sequencing and Target Capture further shows that Target Capture substantially increased mitogenome recovery depth (Supplementary Figure 6). Among sample-reference combinations detected by both methods, Target Capture consistently increased both reference breadth and mean coverage. Median fold enrichment ranged from 21-fold to 56-fold for reference breadth (Supplementary Figure 6a) and from 32-fold to 112-fold for mean coverage (Supplementary Figure 6b), depending on the taxon.

In the separate 55-reference mapping analysis, well 504 yielded the highest individual sample-reference mean coverage, reaching 6.32x against the horse mitochondrial reference. In this well, maximum mean coverages against the human, pig, cattle, dog, and chicken references were 2.73x, 1.40x, 1.25x, 0.50x, and 0.25x, respectively. Well 513 also yielded substantial coverage, with maxima of 3.12x against the pig reference, 1.82x against horse, 1.63x against human, 0.97x against dog, 0.93x against cattle, and 0.40x against chicken. These coverage summaries describe recovery against individual references, rather than the number of genera passing Quicksand’s post-classification filters; the latter was highest in well 513. Well 515 showed lower coverage, with maxima of 0.64x against human, 0.58x against cattle, and 0.26x against pig.

## Notes

### Competing Interest Statement

The authors have declared no competing interest.

